# TRPV1-expressing neurons are dispensable for photophobia: evidence from a novel mouse light sensitivity assay

**DOI:** 10.64898/2026.09.16.752141

**Authors:** Jaewon Sim, Alex D. Chapman, Gwendolyn R. Urbain, Geoffroy Laumet

## Abstract

**Background:** Light hypersensitivity, or photophobia, affects nearly 80% of people with migraine and is the most prevalent symptom after headache. Despite its high prevalence, the neural pathways underlying migraine-associated light hypersensitivity remain poorly understood, in part due to the lack of behavioral assays that directly assess light sensitivity in mice. Consequently, whether nociceptors contribute to the development of photophobia remains unclear.

**Methods:** We established a novel light sensitivity assay (LSA) to quantify the response to light in mice by measuring facial grimacing. The assay was applied to evaluate light hypersensitivity elicited by three established migraine triggers: calcitonin gene-related peptide (CGRP), nitric oxide donor (sodium nitroprusside), and repeated stress. To determine the contribution of TRPV1⁺ nociceptors, mice were treated with resiniferatoxin (RTX) to ablate these neurons.

**Results:** The LSA detected light hypersensitivity following administration of CGRP, sodium nitroprusside, and repeated stress in wild-type (WT) mice. Sex differences were not observed in CGRP- or stress-induced light hypersensitivity. Furthermore, ablation of TRPV1⁺ nociceptors did not reduce light hypersensitivity induced by migraine triggers.

**Conclusions:** These findings present the LSA as a robust, reproducible, and easy to implement behavioral assay for measuring light sensitivity in preclinical migraine models. Our results further demonstrate that TRPV1⁺ nociceptors are dispensable for the development of migraine-associated light hypersensitivity. This assay provides a useful method for investigating the mechanisms underlying light sensitivity and for evaluating potential therapeutic strategies.

## Background

Migraine is a highly prevalent and disabling neurological disorder characterized by recurrent headache attacks and a combination of sensory symptoms such as light hypersensitivity and periorbital hypersensitivity. Light hypersensitivity, or photophobia, is light-induced discomfort that can trigger or exacerbate headaches. Light hypersensitivity is reported by approximately 80% of individuals with migraine and is the most common symptom aside from headaches^1–5^. Despite its clinical importance, the neural mechanisms underlying migraine-associated photophobia remain incompletely understood.

Light is detected by retinal photoreceptors and intrinsically photosensitive retinal ganglion cells, which relay signals to multiple brain regions involved in visual processing and sensory integration. Retinal inputs engage thalamic nuclei that also receive convergent nociceptive information from the trigeminal system, providing a potential anatomical substrate through which light may aggravate headache pain^6–8^. Additional projections from the retina to hypothalamic and brainstem structures have also been implicated in migraine-associated light sensitivity^9–11^. Together, these observations support the existence of neural circuits capable of linking light perception to pain processing; however, the precise cellular mechanisms responsible for photophobia remain undetermined. One unresolved question is whether peripheral nociceptors are required for the development or expression of photophobia. Sensory neurons expressing transient receptor potential vanilloid 1 (TRPV1) are commonly defined as nociceptors^12^. They have been implicated in migraine pain pathophysiology through their responsiveness to inflammatory mediators and their capacity to release calcitonin gene-related peptide (CGRP), triggering neurogenic inflammation^13–15^. Nevertheless, whether TRPV1-expressing neurons are necessary for light aversion remains unclear. Determining the involvement of nociceptive pathways in photophobia is critical for understanding how light sensory and pain circuits interact during migraine and identifying therapeutic targets capable of alleviating migraine associated photophobia.

Neural mechanisms underlying photophobia can be deciphered using animal models. However, unlike mechanical or thermal hypersensitivity, which can be readily quantified using standardized behavioral assays, the assessment of light sensitivity in animals remains challenging. The most frequently employed method is the light/dark box assay, in which rodents freely move between illuminated and dark compartments, and the time spent in each compartment is used as an indicator of photophobic behavior^16–21^. However, this method is inherently influenced by locomotor activity, anxiety-like behavior, exploratory drive, and motivational factors, making it difficult to specifically isolate light sensitivity. Alternative approaches, such as measuring blink or pupillary reflexes, or employing keypoint-tracking frameworks coupled with complex neural network analyses, often require advanced computational expertise and sophisticated data-processing procedures^22–25^. Building on the well-validated Mouse Grimace Scale (MGS)^26–28^, we developed a simple, cost-effective, and reproducible behavioral assay that does not require extensive training, complex pipelines of analysis, or specialized equipment. Importantly, this assay is independent of the animal’s locomotor and exploratory activities, allowing for a more direct assessment of light sensitivity.

In the present study, we developed a novel light sensitivity assay (LSA) by adapting the MGS to score the response to light and used this approach to investigate the contribution of TRPV1-expressing sensory neurons to light hypersensitivity in response to migraine triggers. Our findings reveal that these neurons play a limited role in light hypersensitivity, suggesting that, in mouse models, photophobia can arise largely independently of classical nociceptive pathways.

## Methods

### Animals

All experiments were approved by the Institutional Animal Care and Use Committee at Michigan State University (MSU) and in accordance with U.S. National Institute of Health (NIH) guidelines. C57BL/6J (WT, #000664) mice were obtained from Jackson Laboratory (Bar Harbor, USA) and bred at the MSU animal facility. Mice had *ad libitum* access to food and water and were housed under a 12-h light/12-h dark cycle under the condition of controlled humidity and temperature. Female and male mice were 8–10 weeks of age at the time of experimentation.

### Drug administration

Resiniferatoxin (RTX) (Alomone Labs, R-400) was administered subcutaneously in the flank of 4-week-old WT mice on three consecutive days at doses of 30 mg/kg, 70 mg/kg, or 100 mg/kg. Control mice received vehicle solution (PBS 1X + Tween 80 + DMSO). Both groups rested for four weeks, and a hot plate test (52°C) was performed to confirm the ablation as previously described^26,29,30^. Rat calcitonin gene-related peptide (CGRP) (GenScript, RP11095-0.5), dissolved in 1X PBS, was intraperitoneally injected (0.01 mg/kg). Nitric oxide (NO) donor sodium nitroprusside (SNP) (1 mg/kg) (Millipore-Sigma, #1614501) was dissolved in 1X PBS and administered intraperitoneally to induce migraine in mice^31–33^. Injection volumes were 10 µL/g body weight. Vehicle-treated mice received 1X PBS as a negative control. Mice were tested three hours after injection.

### Restraint stress

Restraint stress was performed following the methods of previous migraine studies^34,35^. Mice were placed in 50 mL centrifuge tubes that had breathing holes drilled. Mice were positioned facing the tapered end of the tube. While crumpled paper towels were added to minimize the movement, adequate space was maintained for normal respiration. Mice were restrained for 2 hours a day for three consecutive days. Restraint stress began between 9:00 – 10:00 AM. Mice were checked every 30 minutes to ensure breathing and appropriate positioning. Restraint stress was performed in a room separate from the animal housing room, while control animals remained in their home cages.

### Light sensitivity assay (LSA) set-up

The following materials were used during LSA: a camcorder with an SD card, acrylic cubes (8.9 × 8.9 × 8.9 cm), a gray paint, an LED lamp (Brightness: 490 lux) partition, disposable puppy pads, a reusable and washable acrylic cube with an open end, two tables of equal height, and a dark-colored solid panel. The outer surface of acrylic cubes was painted gray to optimize visibility on recordings (Figure 1A, B).

**Figure 1.**
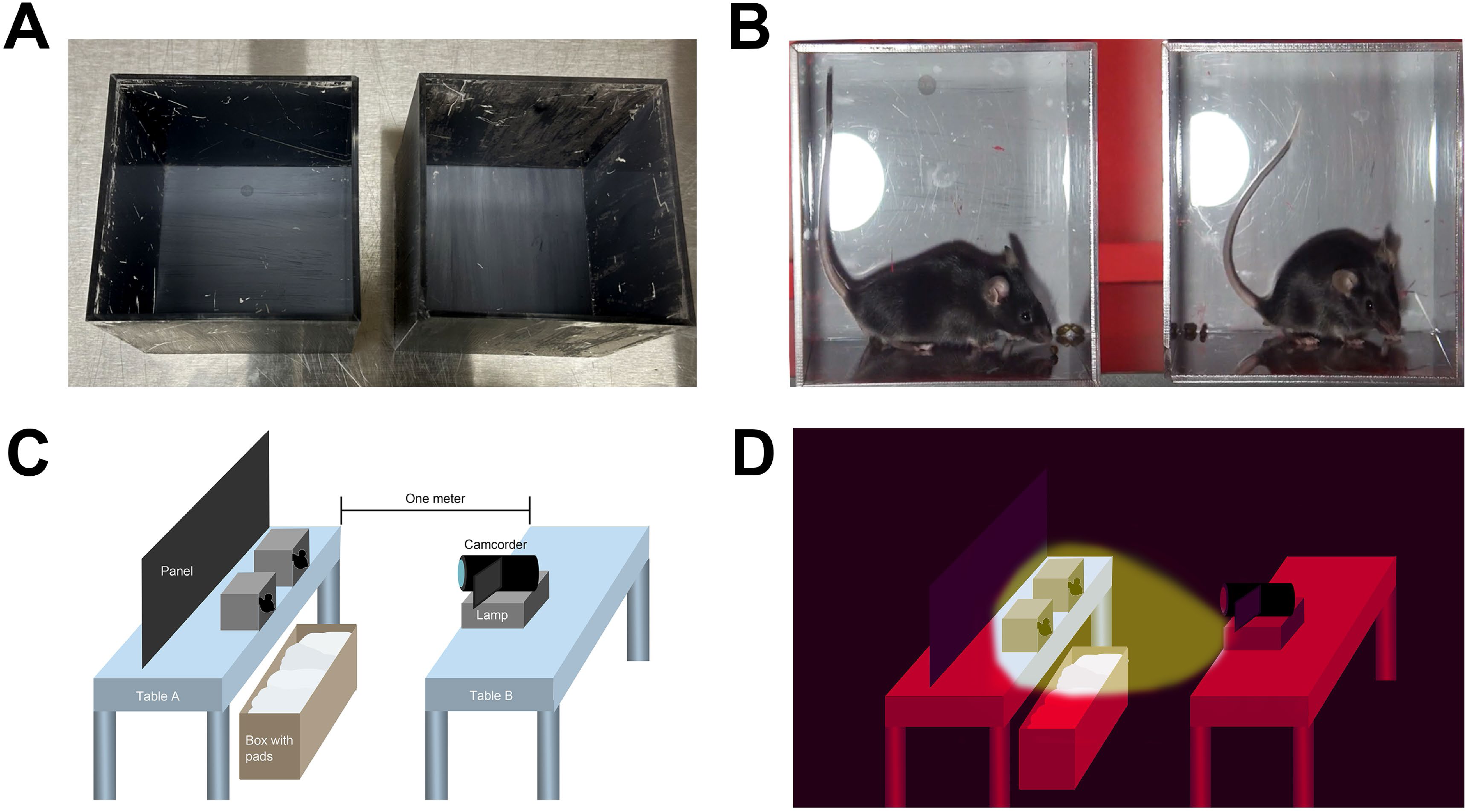
Experiment setup description. (**A**) A painted acrylic cube used for the experiment. Dark gray paint was applied only to the outer surface of the box. (**B**) Cubes with mice inside. (**C**) Illustrations depicting the experimental setup in light and (**D**) dark.

A schematic of the LSA experimental setup is shown in Figure 1C,D. The testing room was equipped with red LED lighting to permit procedures to be performed in darkness. Table A contained two acrylic cubes positioned with their open sides facing the camera and light source. Dark solid panels were placed behind the cubes to minimize light reflection during recording. A padded box was positioned beneath the table to prevent injury if a mouse exited the cube. Table B held the lamp and camcorder, positioned 1 m from the acrylic cubes.

### LSA habituation

The habituation for this assay was spread over three days, not necessarily consecutively. Each day was divided into two phases: habituation to the behavioral room and darkness and habituation to the acrylic cube. During the first 30 minutes (habituation to the behavioral room and darkness), all mouse cages were kept in a dark room illuminated only by red light. After this 30-minute period, habituation to the acrylic cubes began. Mice were habituated to the cubes one cage at a time, approximately 4-5 mice at once. Each mouse was individually placed inside an acrylic cube positioned on a table with the open side facing the edge of the table (Figure 1C). A box overlayed with pads was placed underneath the table to prevent injury or escape if a mouse falls. All procedures were performed under red light. The habituation to the acrylic box phase lasted 15 minutes per group. As habituation progressed, fewer mice attempted to jump or display signs of anxiety inside the cube. Mice were frequently monitored for unusual behavior or falling into the box. Although rare, mice that repeatedly fell were excluded from the experiment. The percentage of mice excluded due to this reason was less than 5% of the total mice across experiments. By the end of the final habituation day, most mice acclimated remaining calm inside the acrylic cube. The habituation procedure was repeated for three days at the same time each day.

### LSA recording

The pre-test was conducted before treatment with migraine triggers, and the post-test was conducted after treatment.

For the first 30 minutes, mouse cages were kept in a dark room. Then, each mouse was placed in an acrylic cube, and the cube was positioned at the edge of the table, as done during habituation. After 5 minutes of acclimatation, the light was turned on and the recording began. The light source was positioned to directly illuminate the mouse (Figure 1D). Two mice were tested at a time. We ensured that mice not currently being tested are not exposed to the light. During the 5-minute video recording period, the experimenter remained in the room as quiet and still as possible. Acrylic cubes were cleaned with water and wiped dry before testing the next round.

The duration of video recording was determined based on a previous study showing that this time frame is sufficient for rodents to exhibit clear place aversion to light stimuli in migraine model^36^. Correspondingly, in humans, exposure to sunlight can trigger a migraine attack within 5 minutes, which aligns with the duration used in our experiment^37^.

### LSA analysis

Before analysis, still images were captured from the recorded videos. Brightness and contrast of video were adjusted using Windows Media Player (brightness: 130; contrast: 110). The first image was captured 30 s after the light was turned on, and additional images were captured every 40–70 s, for a total of five images over the 5-min recording period. Images were captured only when the mouse was stationary and were not captured while the mouse was sniffing, exploring, or grooming. If capturing five images was not possible, fewer images were used. Scoring captured images was adapted from the grimace scoring criteria developed by the Mogil lab (Figure 2)^27^. Eyes, ears, cheeks, and noses were scored on a scale of 0-2, and this scoring is feasible with dark furred C57BL/6 mice (Figure 2). The average grimace score for each mouse was calculated using the images from the same test session, with a maximum score of 2.

**Figure 2.**
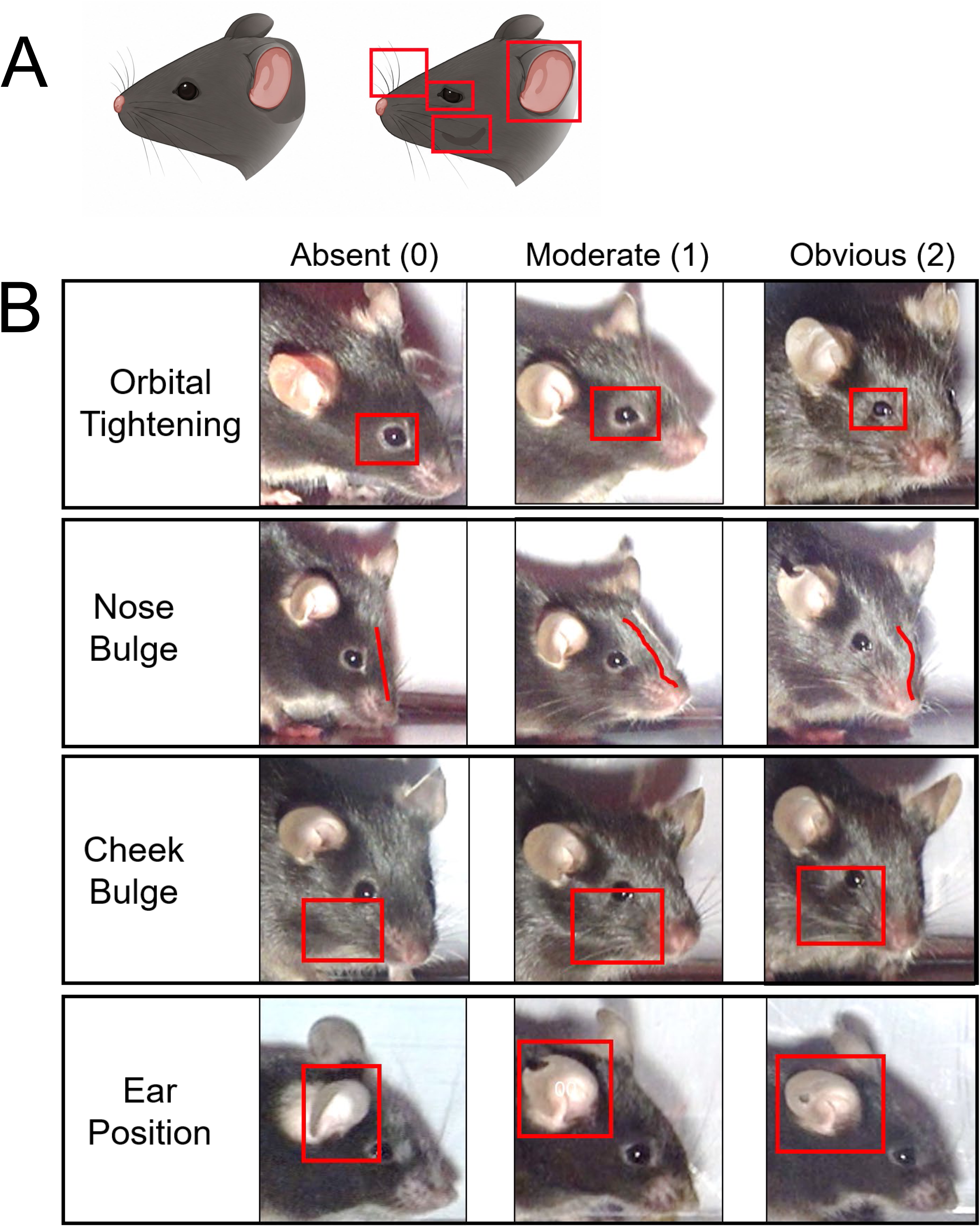
Light Sensitivity Assay (LSA) scoring criteria based on the Mouse Grimace Scale (MGS). (**A**) Schematic illustration of the 4 facial action units scored from recorded videos. (**B**) Representative images illustrating the scoring criteria for each facial action unit, in C57BL/6J mice dark-furred mice, adapted from the Mouse Grimace Scale: orbital tightening, nose bulge, cheek bulge, and ear position. Each feature is scored on a three-point scale as absent (0), moderate (1), or obvious (2). The average score across all facial action units is used to quantify light-evoked grimacing.

### Blinding Procedures

To prevent bias and ensure blinding, a random identification was assigned to each mouse and an experimenter blinded to the condition scored the images.

### Facial mechanical sensitivity

Facial mechanical sensitivity was measured using the von Frey filaments applied to the forehead region of the mice as previously described^35,38^. During four to five habituation sessions, mice were confined in a transparent acrylic cube (8.9 × 8.9 × 8.9 cm) together with a Choice 4-oz cup containing a food pellet. Each session lasted 2 hours, and only one session was performed per day. Mice that adapted to the cup proceeded to the baseline measurement. Baseline measurements were repeated four times. Starting with a 0.07 g filament, the Dixon up-down method was used to calculate the withdrawal threshold. A response was recorded when the mouse swiped the filament after the filament had been sufficiently bent. Mice with a final baseline facial withdrawal threshold greater than 0.47 g but less than 0.7 g were included in the subsequent test experiments monitoring facial mechanical sensitivity overtime.

### Hot plate test

Thermal nociception was assessed using the hot plate test. Mice were individually placed on a hot plate maintained at 55°C, and the latency to the nocifensive responses, defined as hindpaw licking or jumping, were recorded. A cutoff time of 60 s was used to prevent tissue injury. The experimenter was blinded to treatment group.

### Enzyme-Linked Immunosorbent Assay (ELISA)

We followed an established protocol, briefly, mice were euthanized using carbon dioxide. Kidney and trigeminal ganglia were collected, and the tissues were weighed, homogenized in 0.7 mL of 1 M acetic acid, boiled for 20min (95 °C), and centrifuged at 20,000 g for 60 min at room temperature. The supernatants were collected, and the pellets were washed with 0.4 mL of 0.1 M acetic acid, then centrifuged again at 20,000g for 20 min. The supernatants from the second centrifugation were added to the first one. Then, each sample was mixed with 1 mL of Buffer A (1% Trifluoroacetic acid), centrifuged at 12,000 g for 20 min (4 °C). The CGRP was extracted with Buffer B (60% acetonitrile/1% Trifluoroacetic acid, 1ml, 3x) using peptides were extracted with Oasis HLB 3 cc Vac 60 mg tubes (WAT094226, Waters, Colorado Springs, CO). The elution was dried out using a speed vacuum for 5 hours (50 °C for 3 hours, room temperature for 2 hours, pressure 0.5psi) (Savant™ SpeedVac™ SPD120, Thermo Scientific™, USA). The extracted pellets were reconstructed with 0.25 mL of EIA buffer from the CGRP ELISA kit (item no. 589001, Cayman Chemicals, Ann Arbor, MI). CGRP measurement was performed following the manufacturer’s instructions. To eliminate any intra-assay variance, all samples were run on a single 96-well ELISA plate. The concentrations were analyzed with GainData Software and normalized by tissue weight.

### Immunofluorescence imaging for trigeminal ganglia and meninges

Mice were perfused with ice-cold PBS and 4% formaldehyde, and trigeminal ganglia (TG) and meninges were harvested and placed into 4% formaldehyde for fixation and cryoprotected. The TG tissue was cut into 20 µm thick sections on a Leica CM3050 S cryostat. TG sections were incubated with guinea pig anti-TRPV1 (1:200, PA1-29770, Invitrogen, Carlsbad CA), rabbit anti-CGRP (1:500, 24112, Immunostar, Hudson, WI), and anti-mouse NeuN (1:500, 66836-1-1g, Proteintech, Rosemont, IL). The sections were then incubated with Alexa fluor 488 anti-guinea pig (1:300, A11073, Invitrogen, Carlsbad CA), Alexa fluor 568 anti-rabbit (1:300, A11011, Invitrogen, Carlsbad, CA), and Alexa fluor 647 anti-mouse (1:300, A21236, Invitrogen, Carlsbad, CA). Omitting primary antibodies resulted in the absence of staining.

The meninges were peeled off the skull in ice cold PBS. Free floating meninges were incubated with hamster anti-CD31 (1:200, MAB1398Z, Sigma, St. Louis, MO) and rabbit anti-CGRP (1:500, 24112, Immunostar, Hudson, WI). The samples were then incubated with Alexa fluor 488 anti-hamster (1:300, A78963, Invitrogen, Carlsbad CA) and Alexa fluor 568 anti-rabbit (1:300, A11011, Invitrogen, Carlsbad, CA). Omitting primary antibodies resulted in the absence of staining. Sections were visualized on a Leica Stellaris microscope.

### Experimenters

Three independent experimenters, all blinded, were able to perform the LSA and obtained consistent data in both sexes. One caveat was that we did not test whether male experimenters would obtain similar data.

### Illustrations

BioRender, OpenAI, and Adobe Photoshop were used for illustrations.

### Statistics

Data were presented as mean ± error of the mean (SEM). The statistical test for each analysis is described in figure legends. Statistical differences are illustrated as follows ns, not significant, *p < 0.05, **p < 0.01, ***p < 0.001, ****p < 0.001. Test and statistical differences and graphics were made with GraphPad Prism 10.

## Results

### The Light Sensitivity Assay detects photophobia induced by migraine triggers

We aimed to develop a simple, cost-effective, and reproducible test of light sensitivity in mice that can be easily adopted across laboratories and is not confounded by motivation, anxiety-like behavior, or locomotor function. First, we assessed whether migraine triggers known to induce migraine-like behaviors in mice affect the behavioral response measured by our LSA. We determined the effects of three migraine triggers; calcitonin gene-related peptide (CGRP)^17,39^, sodium nitroprusside (SNP)^31,40^, and stress^35,41^.

First, we performed the light-sensitivity assay (LSA) in vehicle- or CGRP-treated mice (Figure 3A, B). Vehicle-treated mice showed similar grimacing scores in response to light before and after treatment. In contrast, CGRP-treated mice exhibited more than a twofold increase in grimace scores relative to their pre-test baseline. A significant increase in grimace scores relative to baseline was considered indicative of light hypersensitivity.

**Figure 3.**
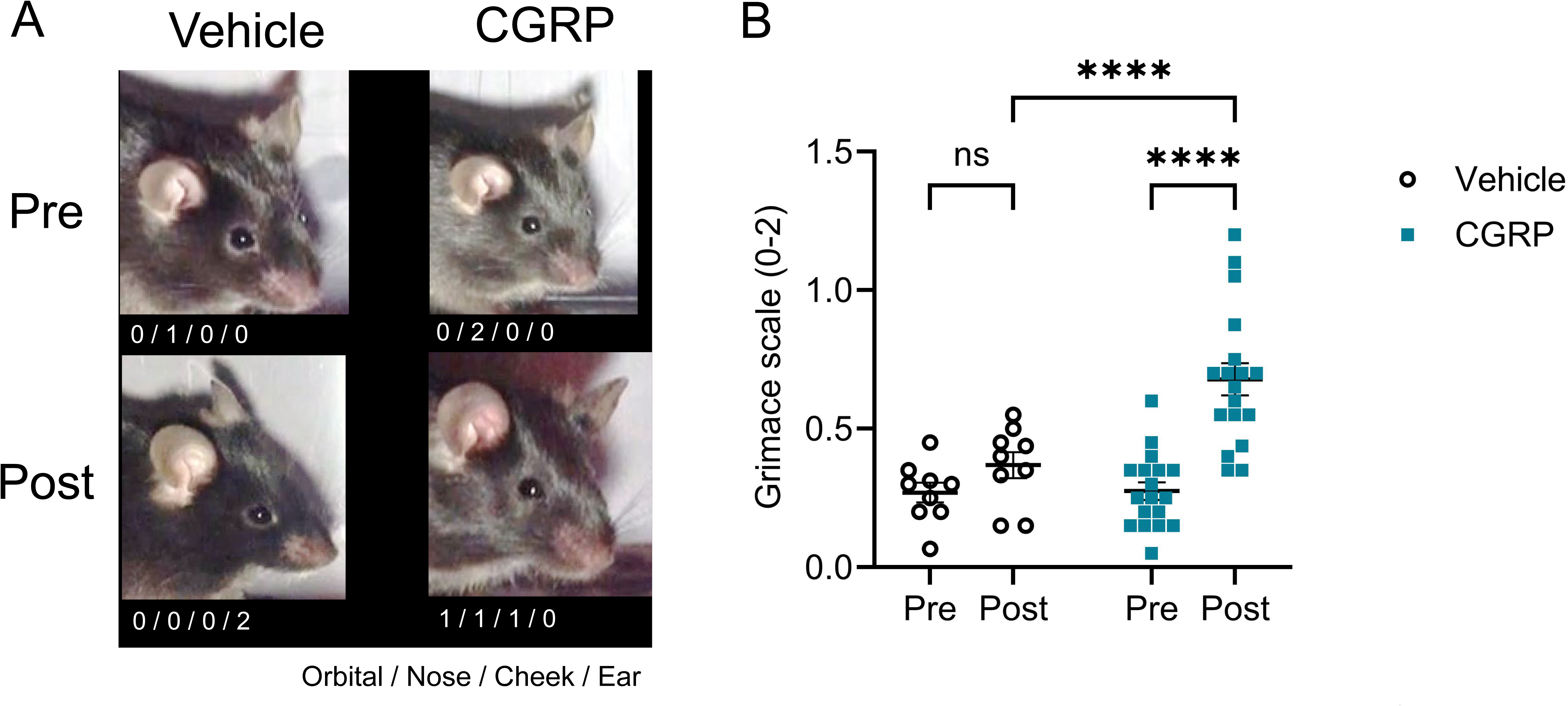
CGRP administration increases light sensitivity. Calcitonin gene-related peptide (CGRP) was administered intraperitoneally (0.01 mg/kg). The vehicle group received 1X PBS (100 μL/10 g body weight). LSA was conducted 3 hours after administration. (**A**) Representative photos of WT mice during the pre-test session (before administration) and post-test session (after administration). (**B**) CGRP treatment increased grimace scores under bright light (indicating increased light sensitivity) compared with vehicle treatment in WT mice. (Vehicle: 4 males, 5 females; CGRP: 8 males, 10 females.) Repeated-measures two-way ANOVA was used.

Similar to CGRP treatment, SNP-treated mice displayed significantly elevated light hypersensitivity compared with their pre-treatment baseline (Figure 4). Across two clinically relevant pharmacological migraine triggers, the LSA consistently detected light hypersensitivity, demonstrating its ability to capture migraine-associated responses to light.

**Figure 4.**
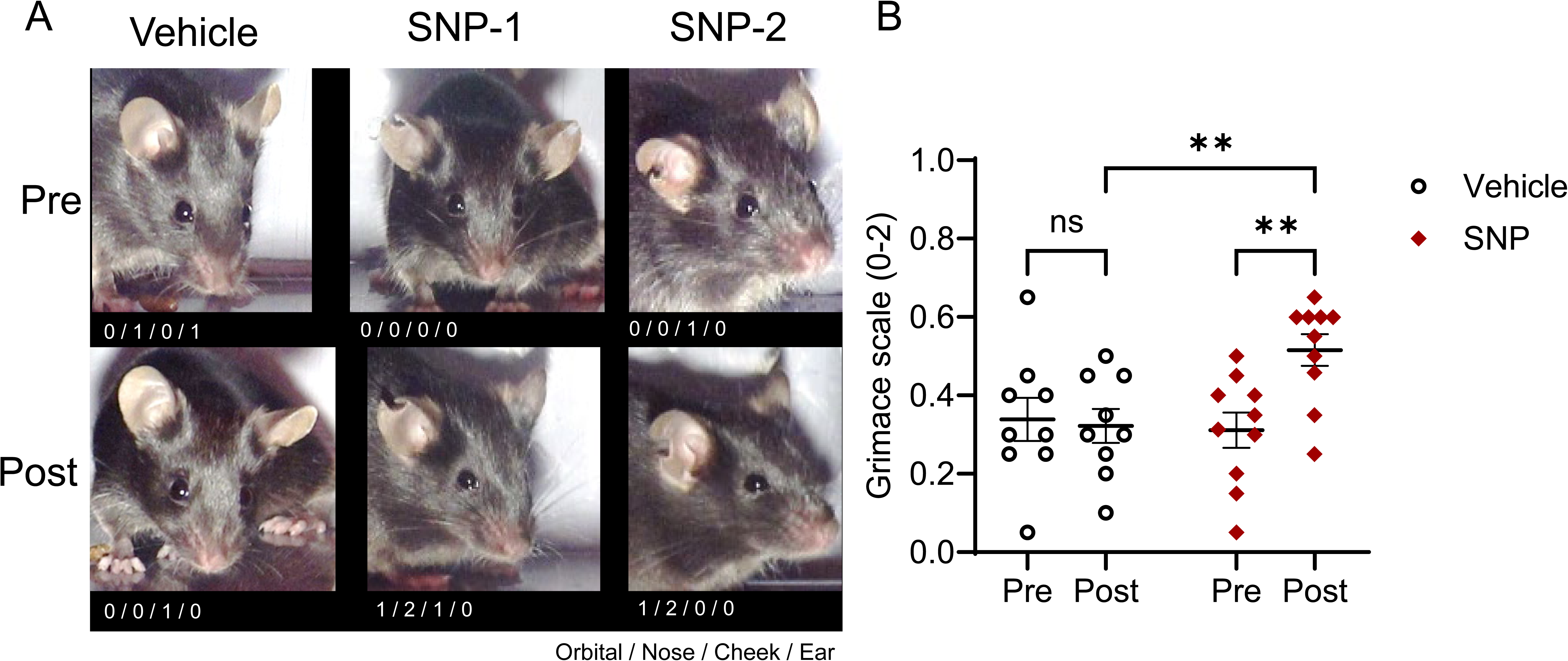
Nitric oxide donor increases light sensitivity. Sodium nitroprusside (SNP) was administered intraperitoneally (1 mg/kg). The vehicle group received 1X PBS (100 μL/10 g body weight). LSA was conducted 3 hours after administration. (**A**) Representative photos of WT mice during the pre-test session (before administration) and post-test session (after administration). (**B**) SNP treatment increased light sensitivity compared with vehicle treatment in WT mice. (Vehicle: 9 females; SNP: 10 females). Repeated-measures two-way ANOVA was used.

We next investigated whether our LSA could also detect light hypersensitivity in a more physiological migraine model that is not induced by pharmacological manipulation. Stress is the most common trigger of migraine in humans^4,41,42^, and previous studies have shown that restraint or sound stress promotes migraine-like behaviors in mice^35,43,44^. Using a validated restraint stress paradigm^35,45,46^, we assessed light-induced grimacing in stressed animals. While non-stressed control mice showed no change in grimace scores, mice subjected to restraint stress exhibited significantly increased grimacing in response to light exposure (Figure 5). These results demonstrate that the LSA also detects light hypersensitivity in a physiologically relevant stress model of migraine.

**Figure 5.**
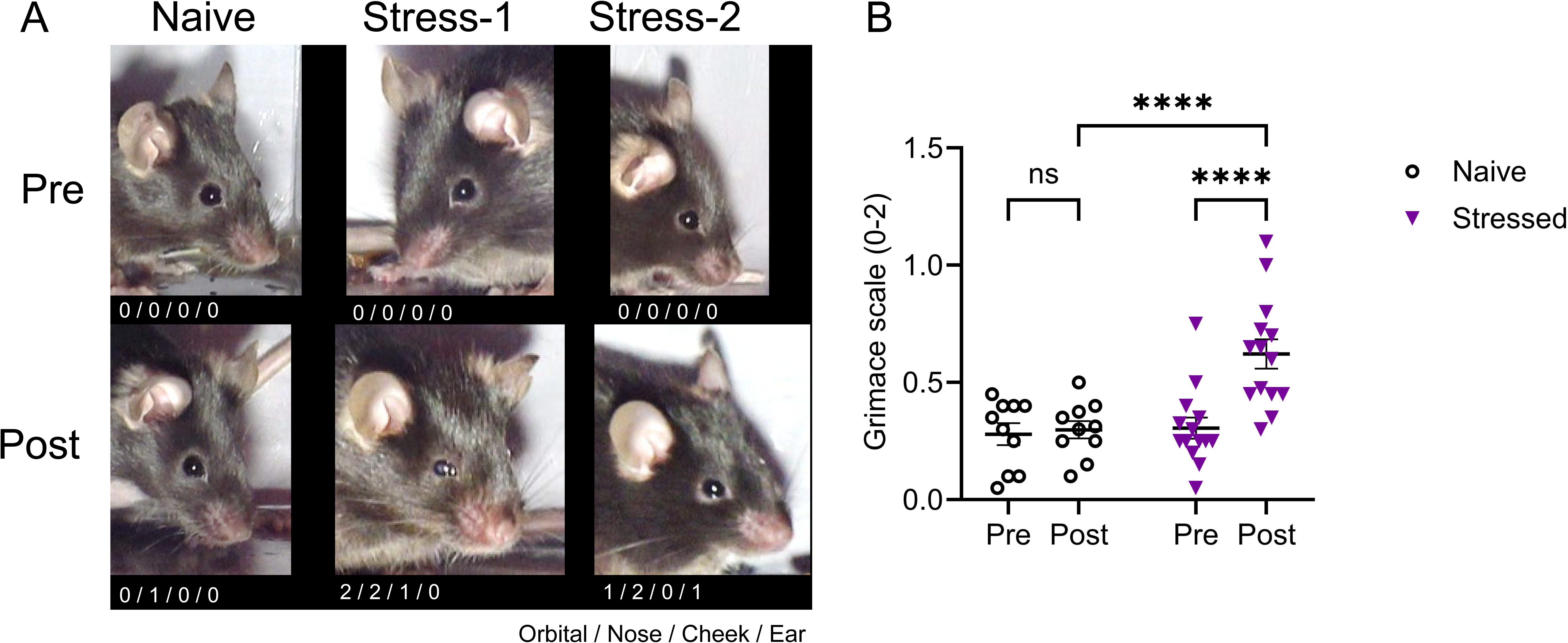
Restraint stress increases light sensitivity. Restraint stress was conducted 2 hours a day for three consecutive days. LSA was performed 48 hours after last session of stress. (**A**) Representative photos of WT mice during the pre-test session (before stress) and post-test session (48 hours after stress). (**B**) Restraint stress increased light sensitivity compared with naive WT mice. (Naive: 2 males, 8 females; Stressed: 6 males, 8 females). Repeated-measures two-way ANOVA was used.

Assessment of facial mechanical sensitivity is commonly used to evaluate migraine-like behaviors in rodents. We therefore examined whether light hypersensitivity and facial mechanical hypersensitivity correlate (Figure 6). Lower facial withdrawal thresholds, indicative of greater mechanical hypersensitivity, correlated with higher grimacing scores. This correlation supports the LSA as a behaviorally relevant readout of migraine-associated sensory hypersensitivity.

**Figure 6.**
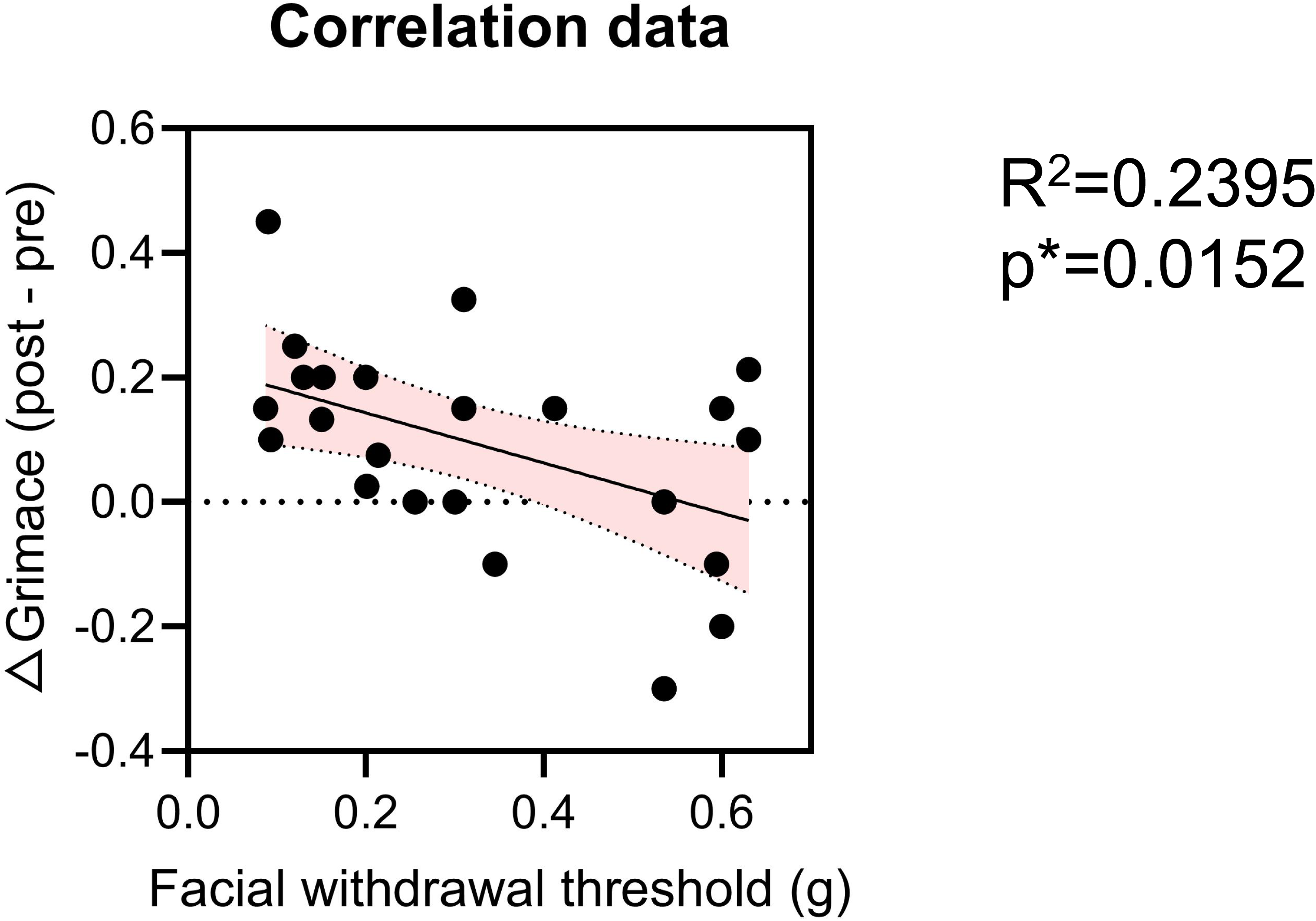
Light sensitivity correlates with facial mechanical sensitivity. X-axis: Facial withdrawal threshold measured 48 hours after restraint stress. Y-axis: Change in grimace score (post-test minus pre-test). Simple linear regression and Pearson’s correlation tests were performed. r = −0.4894; R² = 0.2395; P = 0.0152 (13 females, 11 males).

### Influence of light intensity and sex on the Light Sensitivity Assay

To further characterize the sensitivity of the LSA, we next examined whether increasing light intensity produced graded behavioral responses. Migraine-like hypersensitivity was induced by CGRP administration or restraint stress as described above. Post-tests were performed in the dark, under ambient light (270 lux), or under bright light (490 lux). In the dark, neither CGRP nor restraint stress increased grimace scores relative to pre-test baseline (Figure 7A, C). In contrast, light exposure elicited a significant increase in grimace scores in a light intensity-dependent manner, with responses observed under ambient light and further enhanced under bright light (Figure 7A, C). This intensity-dependent response was observed in both the CGRP and restraint stress models, demonstrating that the LSA is sensitive to the intensity of the light stimulus (Figure 7B, D).

**Figure 7.**
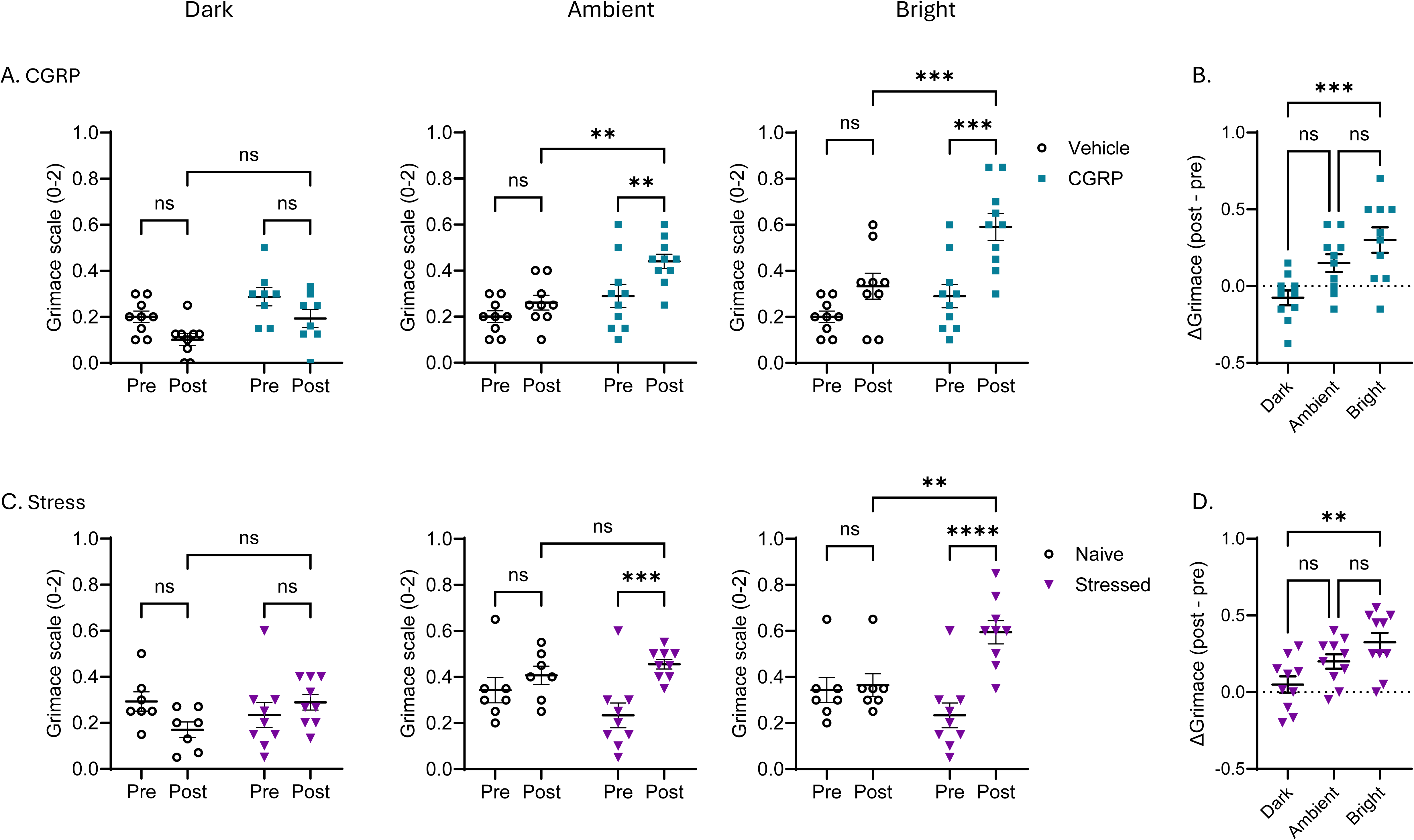
Increasing light intensity induces graded behavioral responses in the LSA. Grimacing was assessed under the dark, ambient light (270 lux), or under bright light (490 lux). (**A**) Calcitonin gene-related peptide (CGRP) was administered intraperitoneally (0.01 mg/kg). The vehicle group received 1X PBS (100 μL/10 g body weight). (**B**) Change in grimace score (post-test minus pre-test) under 3 light conditions. (**C**) Restraint stress was conducted 2 hours a day for three consecutive days. LSA was performed 48 hours after last session of stress. (**D**) Change in grimace score (post-test minus pre-test) under 3 light conditions. <u>Figure 7. Increasing light intensity induces graded behavioral responses in the LSA.</u> Grimacing was assessed in the dark, under ambient light (270 lux), or under bright light (490 lux). (**A**) Calcitonin gene-related peptide (CGRP; 0.01 mg/kg) or vehicle (1× PBS; 100 μL/10 g body weight) was administered intraperitoneally, and grimace scores were assessed under the three light conditions. (**B**) Change in grimace score (post-test minus pre-test) following CGRP or vehicle administration as a function of light intensity. (**C**) Mice were subjected to restraint stress for 2 h/day for three consecutive days, and the LSA was performed 48 h after the final stress session under the three light conditions. (**D**) Change in grimace score (post-test minus pre-test) following stress or control conditions as a function of light intensity.

Sex differences are well documented in both migraine patients and preclinical models^47–49^. Both CGRP administration and restraint stress significantly increased grimacing in female and male mice, with no significant difference in the magnitude of the response between sexes (Figure 8A, B). Thus, under these experimental conditions, our LSA did not reveal sex-dependent differences in light hypersensitivity.

**Figure 8.**
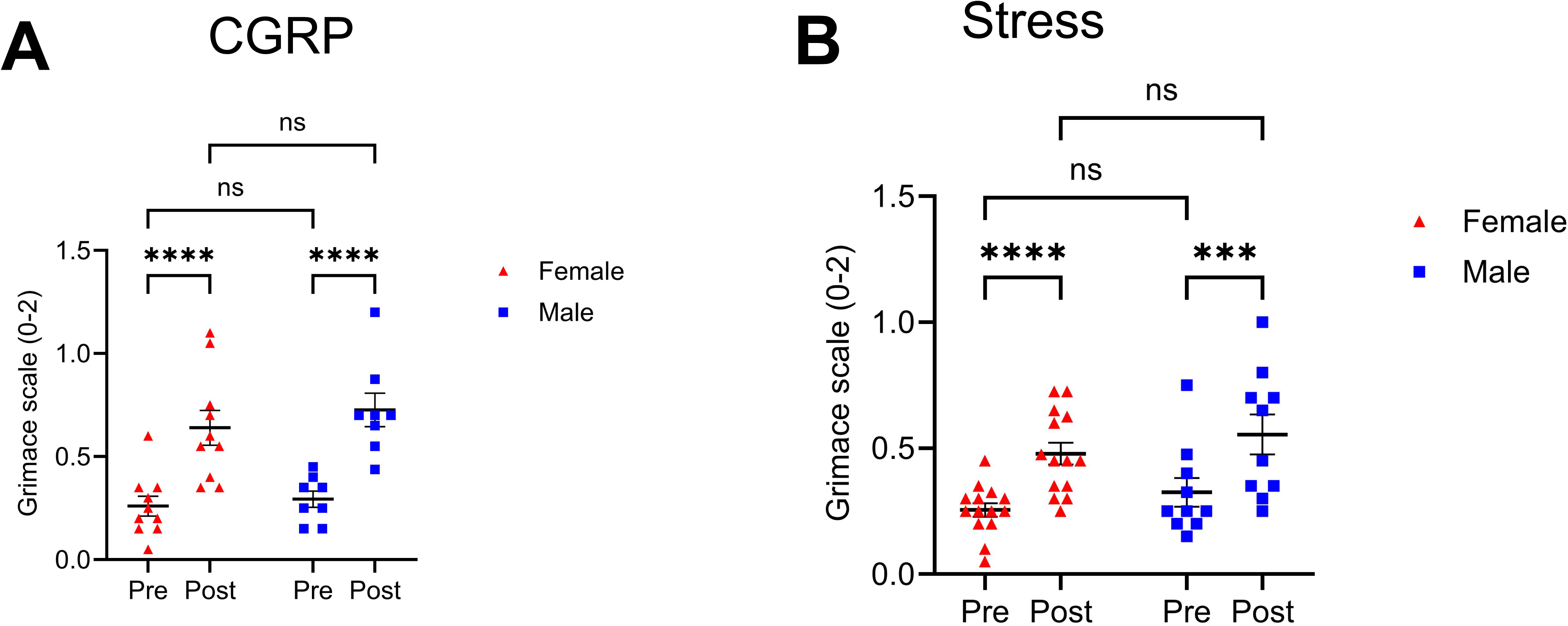
Sex difference in light hypersensitivity was not detected using the LSA. (**A**) Calcitonin gene-related peptide (CGRP) was administered intraperitoneally (0.01 mg/kg). The vehicle group received saline (100 μL/10 g body weight), n=10 females, 8 males. (**B**) Restraint stress was conducted 2 hours a day for three consecutive days. LSA was performed 48 hours after last session of stress, n= 14 females, 10 males. (**A, B**) Repeated-measures two-way ANOVA was used.

### Role of TRPV1+ nociceptors in light hypersensitivity

Having established the validity of the LSA, we investigated the contribution of TRPV1^+^ nociceptors to photophobia. TRPV1^+^ nociceptors were ablated with RTX following a well-established protocol^26,29,50^. RTX treatment markedly reduced thermal sensitivity and CGRP protein levels in end-organ (Figure 9A), thereby confirming the ablation of nociceptors. We then challenged RTX- or vehicle-treated mice with CGRP or restraint stress and assessed light sensitivity. As expected, vehicle-treated mice exhibited increased grimacing following CGRP administration. RTX-treated mice showed similar light hypersensitivity after CGRP treatment (Figure 9B). Regardless of RTX treatment, stressed mice displayed comparable light hypersensitivity (Figure 9C). Collectively, these findings indicate that TRPV1^+^ neuron ablation does not prevent the development of light hypersensitivity, suggesting that TRPV1^+^ nociceptors may not be required for photophobia-like behaviors.

**Figure 9.**
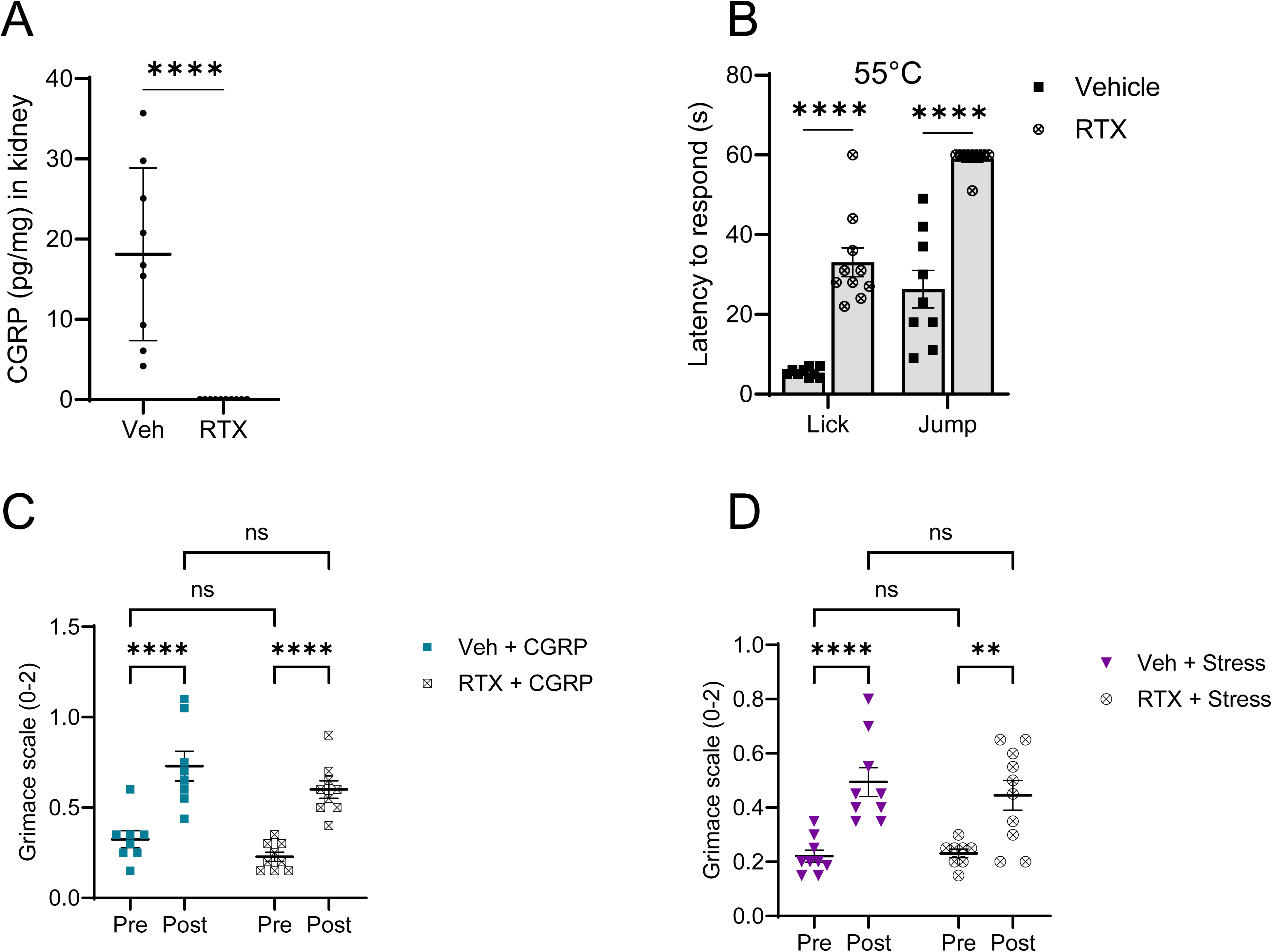
RTX did not prevent light hypersensitivity induced by CGRP or stress. Resiniferatoxin (RTX) was injected to ablate TRPV1^+^ nociceptors. (**A**) CGRP ELISA from kidney samples confirmed the ablation of TRPV1+ nociceptors (unpaired T-test). RTX-treated mice showed reduced response to noxious heat (55°C) (Two-way ANOVA with Tukey’s correction). (**B**) Calcitonin gene-related peptide (CGRP) was administered intraperitoneally (0.01 mg/kg). The vehicle group received saline (100 μL/10 g body weight). (Vehicle: 5 females, 3 males; RTX: 5 females, 4 females.) (**C**) Restraint stress was conducted 2 hours a day for three consecutive days. LSA was performed 48 hours after last session of stress. (Vehicle: n= 5 females, 4 males; RTX: n= 5 females, 5 males.) (B, C) Repeated-measures two-way ANOVA was used.

## Discussion

Light hypersensitivity, or photophobia, affects the majority of patients with migraine and can trigger or exacerbate headache attacks^41,42,51^. Despite its prevalence and clinical significance, the neural mechanisms underlying light hypersensitivity remain poorly understood. To facilitate the study of photophobia in animal models, we developed and validated a novel behavioral test, termed the Light Sensitivity Assay (LSA). The LSA detected light hypersensitivity induced by multiple migraine triggers, including CGRP, a nitric oxide donor, and repeated restraint stress. In addition, ablation of TRPV1^+^ nociceptors did not alter CGRP- or stress-induced light hypersensitivity compared with the control group.

The absence of a requirement for advanced and expensive instrumentation or specialized techniques represents a major advantage of this assay. A standard video camera and a lamp providing appropriate illumination are the primary tools needed for the experiment. Grimacing is scored manually based on the well-established MGS protocol or automatically with analysis software^27,52^. Another advantage of the LSA is that it requires only 5 minutes of recording, thereby reducing the experimental time burden.

Conventional behavioral assays evaluate photophobia indirectly by measuring locomotor responses to light, typically by quantifying the time animals spend in light and dark chambers^16–20,53^. These assays may be confounded by alterations in locomotor and exploratory activity. Notably, several studies have reported impaired locomotor^54–57^ and exploratory^38,58^ behaviors in migraine animal models, suggesting potential challenges in experimental design and data interpretation. In contrast, the LSA directly measures grimacing responses to light exposure without requiring voluntary locomotion. Therefore, this assay provides a simplified, direct, and translationally relevant method for assessing one of the most common and bothersome symptoms of migraine in animal models.

A unique feature of the LSA is its sensitivity to light intensity. Grimacing was absent in the dark and increased in a light intensity-dependent manner, irrespective of the migraine trigger. Thus, the LSA captures a specific behavioral response to light rather than spontaneous pain, providing a direct readout of migraine-associated light hypersensitivity.

One potential limitation of the LSA is that grimacing, particularly eye closure under light, may represent a natural protective response rather than an indicator of migraine. However, we did not observe changes in eye opening in vehicle-treated and control mice. Additionally, a recent study showed that blink frequency did not differ between darkness and light exposure in healthy mice^61^.

Using LSA, we showed that ablation of TRPV1⁺ nociceptors failed to prevent light hypersensitivity. These seemingly contradictory findings suggest that the CGRP driving light hypersensitivity is not derived from TRPV1⁺ sensory neurons. One potential source of CGRP is the parabrachial nucleus in the brainstem. Substantial evidence indicates that parabrachial neurons produce CGRP^61–63^. Although it remains unclear whether CGRP⁺ parabrachial neurons directly influence the neural circuits responsible for light hypersensitivity, brain-derived CGRP may reach the TG through the cerebrospinal fluid (CSF). A recent study demonstrated that, in a cortical spreading depression (CSD) model of migraine, elevated CGRP in CSF directly activates TG neurons that are in close contact with the CSF^64^.

Light hypersensitivity in the absence of trigeminal nociception has also been reported in humans. Patients with migraine frequently experience photophobia during headache-free periods^73,74^. Similarly, individuals with silent migraine can exhibit photophobia and impaired skin sensitivity without headache or other painful sensations^75,76^. These clinical observations match well with our findings that mice continue to exhibit light hypersensitivity even after ablation of TRPV1^+^nociceptive neurons. Collectively, these findings suggest that light hypersensitivity and trigeminal nociception in migraine may be mediated by distinct mechanisms^70,77^.

Overall, LSA represents an effective, simple, cost-efficient, and rapid behavioral assay for assessing light sensitivity in migraine animal models. Improving behavioral testing in animal models is essential for advancing migraine research, by enabling more precise, reliable, and translationally relevant assessment^78–80^. Additionally, LSA may provide a valuable approach for interrogating the neural mechanisms underlying light sensitivity in migraine, independently of the nociceptive aspects of the disorder. These advances would facilitate the development of novel and effective migraine therapies.

## Declarations

No human participants were involved in this study; therefore, consent for publication was not applicable, and human ethics approval and informed consent were not required. All animal procedures were approved (please see Methods, Animal section).

The data generated during the current study are available from the corresponding author upon reasonable request.

### Artificial Intelligence Statement

OpenAI’s ChatGPT was used solely for grammatical and stylistic editing of the manuscript and for the generation of the graphical abstract. The authors reviewed and approved all AI-assisted content and take full responsibility for the final manuscript.

The authors declare that they have no competing interests.

This work was supported by the United States National Institutes of Health (NIH) R21NS142685, R01AI177305 and R01NS121259 (G.L.) and 2T32-GM142521 (A.D.C.), The Rita Allen Foundation (G.L.), the MSU NDRF (G.L.), Red Cedar Distinguished Professorship (G.L.), the John A. Penner fellowship (J.S.), and the Nat Sci Undergraduate Research scholarship (G.R.U).

### Authors’ contributions

Performing Experiments (JS, ADC, GRU), Conceptualization (JS, GL), Methodology (JS, ADC), Visualization (JS), Analysis (JS, ADC, GRU), Supervision (GL), Writing, review & editing (JS, GL). All authors approved the final version of this manuscript for submission.

## Acknowledgements

We thank G. Dussor for sharing expertise and providing training in the facial von Frey test and mouse grimace scoring. We thank Frances M. Seneriz-Pardo (Summer student BPNP) for technical assistance.

## List of abbreviations

(TRPV1): Transient receptor potential vanilloid 1
(CGRP): Calcitonin gene-related peptide
(MGS): Mouse Grimace Scale
(LSA): Light sensitivity assay
(RTX): Resiniferatoxin
(SNP): Sodium nitroprusside
(TG): Trigeminal ganglion
(CSF): Cerebrospinal fluid
(CSD): Cortical spreading depression

